# Spatial layout of visual specialization is shaped by competing default mode and sensory networks

**DOI:** 10.1101/2025.09.08.674858

**Authors:** Ulysse Klatzmann, Robert Scholz, R. Austin Benn, Francesco Alberti, Victoria Shevchenko, Alexander Holmes, Wei Wei, Carla Pallavicini, Robert Leech, Pierre-Louis Bazin, Sean Froudist-Walsh, Daniel S. Margulies

**Affiliations:** Université de Paris Cité, INCC UMR 8002, CNRS, 75006 Paris, France; Wilhelm Wundt Institute for Psychology, Leipzig University, Leipzig, Germany; Max Planck School of Cognition, Leipzig, Germany; Oxford University Centre for Integrative Neuroimaging (OxCIN), Centre for fMRI of the Brain (FMRIB), Nuffield Department of Clinical Neurosciences, John Radcliffe Hospital, University of Oxford, Oxford OX3 9DU, UK; Université Paris-Saclay, Inria, CEA, Palaiseau, France; Buenos Aires Physics Institute (IFIBA) and National Scientific and Technical Research Council (CONICET), Pabellón I, Ciudad Universitaria (1428), Buenos Aires, Argentina; Institute of Psychiatry, Psychology & Neuroscience, King’s College London, London, UK; Full brain picture Analytics, Leiden, The Netherlands; Bristol Computational Neuroscience Unit, School of Engineering Mathematics and Technology, University of Bristol, Bristol BS8 1UB, UK

**Keywords:** Spatial Component Decomposition, Sparse dictionary learning, Retinotopy, Cortical gradients, Population receptive field, Cortical Influences, Distributed representation, Cortical Competition

## Abstract

Understanding how the brain encodes information started with a map but turned into a maze: paths multiplied; boundaries blurred. Neurons tuned to specific features are not confined to single regions, but distributed across the cortex. Retinotopy, once thought limited to early visual areas, now appears in over 20 cortical regions—from the tip of the occipital cortex to the shore of the lateral frontal cortex. To describe and understand these complex mosaics of functional specialization, we focus on the spatial influences that shape their emergence across the cortical sheet. To this end, we developed Spatial Component Decomposition (SCD), a sparse dictionary learning framework that locates sources of spatial influence without relying on prior assumptions from systems neuroscience. Applied to MRI data capturing retinotopic maps, SCD reveals a dominant linear gradient extending over 60 mm from V1 and covering all the known posterior visual areas. Yet, it also revealed systematic competition from other primary sensory areas and default mode transmodal hubs. These suppressive influences shape the cortical embedding of visual information, even during purely visual tasks. Our results suggest that functional specialization emerges from spatial competition between representational systems—not just from feedforward inputs.

## Introduction

A central challenge in systems neuroscience is to understand how mental representations are organized across the brain, and more specifically across the cortical sheet. Classical lesion studies showed that damage to certain brain regions can disrupt specific perceptual or cognitive functions[1], while recordings at the level of populations and single neurons have revealed striking correlations between neural activity and representational content, from low-level sensory features[2, 3] to high-level semantic categories[4]. Yet carefully studied, many of these representations appear not to be confined to individual areas, but distributed across multiple regions[5, 6, 7]. This has led to the concept of population encoding[8], in which information is embedded in the joint activity of neural ensembles rather than localized in single units. Ultimately, most mental representations are distributed across multiple brain regions, and most brain regions contribute to several functions, reflecting a system that encodes information in an overlapping and redundant way. This view aligns with the fact that individual neurons are inherently noisy and variable[9], making them unreliable holders of stable content. However, when considered collectively, like particles in a fluid or molecules in a gas, neurons give rise to emergent dynamics that are far more structured and robust[10, 11, 12]. From this perspective, the core question shifts: not just where information resides, but how distributed codes are spatially organized and embedded in the geometry of cortical sheet.

Retinotopic encoding provides a well-characterized and biologically grounded case study for addressing this question[13]. Its investigation has expanded our understanding beyond the classical “one region, one representation” view. Not only is retinotopic information distributed, appearing in over 20 cortical areas[14, 15, 16], including parietal and frontal regions[17, 18, 19], but it also varies in degree across these areas. Each region with a retinotopic representation only devotes a fraction of its activity to encoding visual field position[20, 21, 17]. This partial involvement highlights a central challenge: to understand brain architecture, we must quantify the contribution of each representation to a region’s activity. In early visual areas such as V1–V4, retinotopic representations drive a large fraction of regional activity[16, 22]. In many higher-order regions, the contribution is smaller yet consistent and functionally meaningful. This gradient can be captured quantitatively using retinotopic selectivity—the fraction of signal variance explained by a region’s preferred visual field location during an experiment. High-resolution population receptive field (pRF) models and large-scale datasets quantify this selectivity across the cortical surface with considerable accuracy[16, 22], at a spatial resolution of only a few square millimeters—orders of magnitude finer than the hundreds of square millimeters typically covered by cortical areas in classical atlases[23, 24]. This allows us to characterize not just where visual information is represented, but the strength of visual encoding across local cortical populations all over the cortex. This graded, distributed architecture provides a rich substrate for probing the organization of sensory representations, and how the geometry of the cortex constrains it.

Two intuitive mechanisms may explain how the geometry of the cortical sheet drives neuronal activity-one developmental, the other dynamic. Developmentally, gradients of morphogens and transcription factors lay down a structural scaffold that guides cortical differentiation[25], leading to regional differences in microcircuit properties such as receptor density[26], myelination[27, 28], or cell-type distribution[29]. These anatomical gradients create long-lasting biases in local computations, predisposing regions toward particular functional roles. The second mechanism is dynamic: the functional role of a region is partly shaped by the activity it receives. Because most cortical connections are local—their likelihood declines sharply with distance[30]—activity tends to propagate as traveling waves across the cortical sheet[31, 32], while long-range white matter tracts provide shortcuts between distant areas. Building on this principle, it is no surprise that task-evoked activity patterns can be efficiently reconstructed using eigenmodes derived from cortical geometry[33], and incorporating sparse long-range connections further improves this predictive power[34]. In both mechanisms, the spatial influences can be viewed as either: (i) a developmental gradient—such as a morphogen diffusing along the cortical surface from an embryonic anchor point; or (ii) a functional gradient—emerging from the propagation of activity initiated at specific cortical entry points by subcortical projections, such as those from the thalamus or hippocampus. Characterizing and isolating these distinct spatial influences is essential for understanding how cortical geometry shapes functional specialization and for building biologically grounded models of distributed representation[35].

To address this challenge, we introduce Spatial Component Decomposition (SCD): a sparse-dictionary-learning framework[36, 37] that isolates a compact set of spatially localized, biologically interpretable components. SCD models both facilitative influences (regions that contribute to the representation) and suppressive influences (regions that systematically reduce it). These hidden patterns reflect how different cortical systems compete or cooperate to shape representational structure. We apply SCD to retinotopic selectivity to reveal how visual-field information is embedded in cortical geometry. The same framework can be extended to other continuous or distributed features—semantic tuning, motor representations, or task activations—where clearer biological anchoring is equally desirable. Applied to retinotopic selectivity, Spatial Component Decomposition (SCD) identified eight spatially organized components per hemisphere—two facilitative, six suppressive—that together account for 92.3% of the explainable variance. This strikingly compact representation reveals that the complex, cortex-wide distribution of retinotopic selectivity can be reduced to a small set of spatial influences. These components are not arbitrary: they are anchored principally to cooperating and competing primary sensory and transmodal hubs of the Default Mode Network (DMN) regions. Together, these results support a view in which functional specialization arises not just from input-driven tuning, but from spatially structured competition among mental representations.

## Results

### Widespread Retinotopic Gradient Beyond Early Visual Areas

Retinotopic selectivity forms a topographic continuum: highest values cluster in early visual cortex, but measurable selectivity extends far beyond (Figure 1A). Because retinotopic selectivity quantifies the proportion of signal variance explained by visual field position, higher retinotopic selectivity indicates a greater degree of specialization for visual field encoding. This broad spatial profile suggests that retinotopic information fades, rather than disappears, with cortical distance, hinting at a macroscale gradient of functional specialization, consistent with classical models of hierarchical sensory processing [38, 39]. We hypothesized that this gradient could be captured by geodesic distance from primary visual cortex (V1), and tested whether spatial proximity to V1 alone could explain large-scale variation in selectivity. We modeled selectivity as a threshold-linear function of distance from V1 (Figure 1B,C), parameterized by maximum value at V1, a spatial decay constant, and a floor value at distal regions. Models were independently fit on each hemisphere using the full unthresholded data to preserve their spatial detail. Despite its simplicity, this geometric model explained 23.8% for the left hemisphere and 20.2% for the right hemisphere of the variance in heldout subjects. For reference, the variance explainable by the group-average selectivity map was 74.1% and 70.6% for the left and right hemispheres, respectively, representing an upper bound given the intersubject variability present in the data.

**Figure 1.**
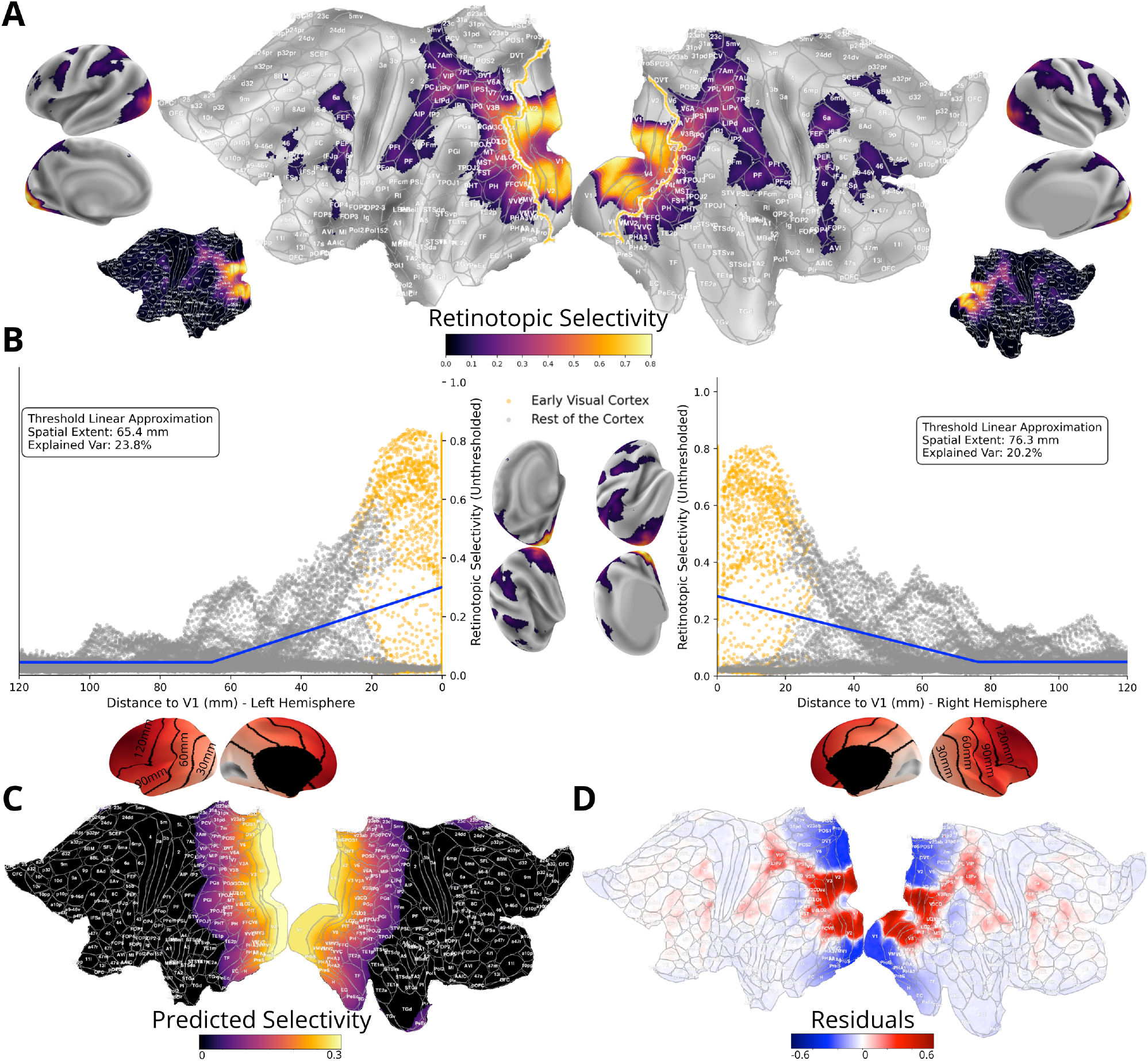
Widespread Retinotopic Gradient Beyond Early Visual Areas. **A**. Retinotopic selectivity across the human cortex. Average selectivity values are shown for both hemispheres on inflated and flattened cortical surfaces, with major anatomical landmarks overlaid. The yellow mask outlines early visual areas (V1-V4) based on the MMP1.0 atlas [24]. A visualization ceiling was applied to enhance contrast, and an unthresholded version is provided in the bottom-left inset. **B**. Scatter plot of unthresholded retinotopic selectivity as a function of geodesic distance from V1. Yellow dots denote vertices within early visual areas (V1–V4); the blue line shows the threshold-linear model fit. **C**. Predicted selectivity from the threshold-linear model, projected onto the flattened cortical surface with anatomical references overlaid. **D**. Model residuals displayed on the same surface, highlighting regions that deviate from purely distance-based prediction.

These results support a central conclusion: retinotopic selectivity declines gradually with cortical distance from V1, extending well beyond early visual areas. While regions like V2–V4 typically fall within 20 mm of V1 (Figure 1A,B), selectivity remained structured over *∼*60 mm of cortex. This slow decay implies that visual encoding is embedded in a continuous cortical gradient, not confined to discrete sensory zones. Critically, this effect cannot be attributed to spatial smoothing or averaging artifacts, which operate over much smaller scales of only a couple of milimeters [40, 41]. The threshold-linear model captured the broad spatial trend but left spatialy structured residuals: some regions systematically over- or under-performed relative to their distance from V1 (Figure 1D). These deviations point to additional spatial influences, motivating a more flexible decomposition of retinotopic organization.

### Repulsive Regions and Facilitating Centers Shape the Retinotopic Map

To capture the structured deviations left unexplained by the threshold-linear model, we extended the previous model, and developed a novel method—Spatial Component Decomposition (SCD). SCD models the retinotopic selectivity map as a sum of spatially localized, signed components on each hemisphere independently (Fig. 2A, Eq. 1). Each component is anchored to a cortical vertex and defined by a linearly decaying influence over the cortical surface, clipped to zero beyond a spatial extent (Eq. 2,3). This allows the model to identify both facilitative and suppressive regions that shape the distribution of retinotopic selectivity.

**Figure 2.**
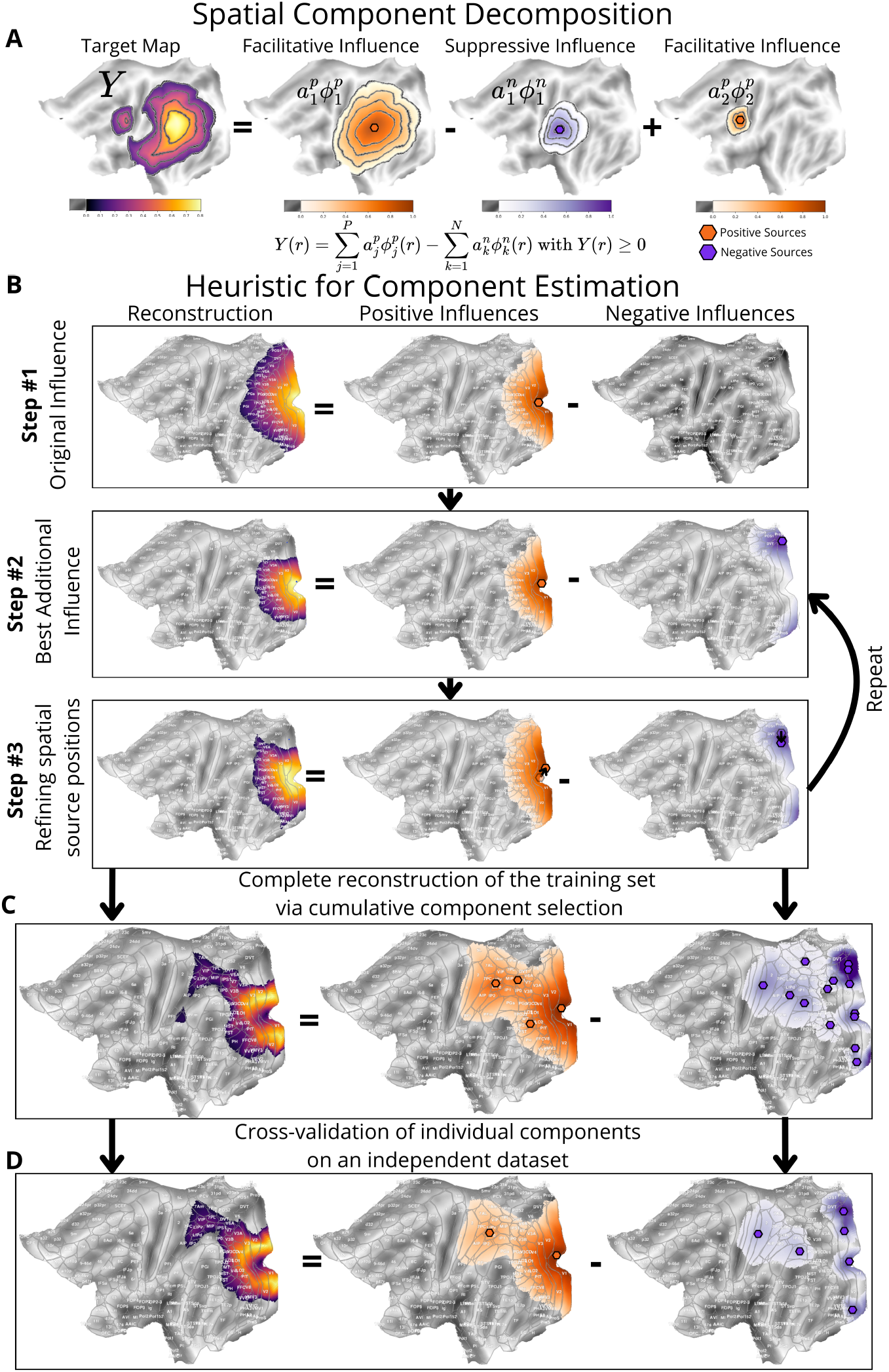
Modeling retinotopic selectivity with spatially localized influences. **A**. Decomposition of an artificially-generated target map into three spatial components (two facilitative and one suppressive), to demonstrate the method. The decomposition formula is displayed, along with the location of the source for each spatial influence. A sulcal background is shown in regions where the value of any map is zero. Yet, all the analysis are performed on the unthresholded maps. **B**. Heuristic used to derive the spatial components. **Step 1:** Identify the initial facilitative influence by selecting the vertex explaining most variance. **Step 2:** Add a new component by choosing an additional vertex that maximizes the variance explained, without altering previously selected positions. **Step 3:** Jointly refine the positions of all existing components locally to improve the fit. Repeat from Step 2. For each step, we show the current reconstruction, as well as the cumulative facilitative and suppressive influences. **C**. Full heuristic applied to the training set. The resulting reconstruction is shown along with the cumulative facilitative and suppressive components and their source locations. **D**. Cross-validation of each component on an independent dataset (NYU retinotopy), components which don’t generalize are droped. Shown are the reconstruction, cumulative facilitative and suppressive components, and the spatial sources of each retained influence.

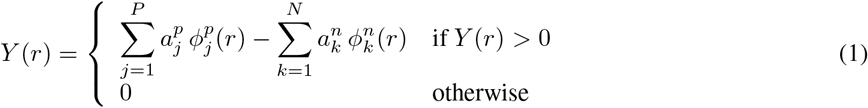

With *Y* denotes the target statistical map for all vertices *r, ϕ*^*p*^ and *ϕ*^*n*^ being respectively the facilitative and suppressive components, and *a*^*p*^ and *a*^*n*^ being the components loadings. Each component is parameterized as a threshold-linear function, as defined in Equations 2 and 3.

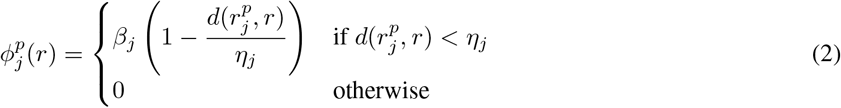

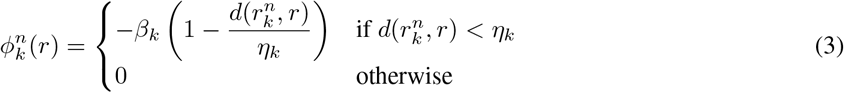

With *d*(*r, r*^*′*^) denoting the geodesic distance between cortical vertices *r* and *r*^*′*^. Here, *η* defines the spatial extent of the component, *β* scales its maximum amplitude, and *r*^*p*^ and *r*^*n*^ specify the cortical locations of the positive and negative sources of influence, respectively.

SCD adapts the sparse dictionary learning framework to the geometry of the cortical surface [36, 37], constraining components to be spatially compact non-negative via a rectifying (ReLU) nonlinearity. Unlike convex regularization methods such as Lasso, sparse dictionary learning presents a non-convex optimization problem with no closed-form solution. As a result, component selection must proceed heuristically, through a stepwise procedure that iteratively builds a dictionary to optimize both sparsity and explanatory power (Figure 2B; see Algorithm 1 for the corresponding pseudocode)). Full details of this optimization strategy are provided in the Methods section. To ensure the robustness of these components, the model was initially fit on the HCP 7T retinotopy dataset [16] (Figure 2C), where the decomposition yielded 18 spatial components for the left hemisphere, and 11 for the right hemisphere. The model was then validated on an independent dataset [22] (Figure 2D), retaining only the components producing a statistically significant improvement in model performance, using a conservative significance threshold of *p* = .0005, Bonferroni-corrected for multiple comparisons. Of the initial components, 8 survived cross-validation in each hemisphere and were used for prediction on the held-out test set (Table 1,2).

**Table 1:** Spatially localized components identified in the left hemisphere using the HCP dataset.

| Seed Vertex (fs_l32) | Atlas (Yeo17 / Glasser) | Influence Type | Extent (mm) | Magnitude |
| --- | --- | --- | --- | --- |
| 23909 | VisCent / V1 | Facilitative | 59.0 | 1.00 |
| 23864 | VisCent / V1 | Suppressive | 19.3 | 0.79 |
| 25760 | VisPeri / V1 | Suppressive | 30.1 | 1.00 |
| 14603 | DorsAttnA / LIPv | Facilitative | 45.4 | 0.44 |
| 16237 | DefaultB / PGs | Suppressive | 30.5 | 0.40 |
| 7436 | SomMotA / Area 1 | Suppressive | 44.6 | 0.38 |
| 25149 | VisPeri / V2 | Suppressive | 18.7 | 0.80 |
| 25272 | VisPeri / V2 | Suppressive | 19.4 | 0.63 |

**Table 2:** Spatially localized components identified in the right hemisphere using the HCP dataset.

| Seed Vertex (fs_l32) | Atlas (Yeo17 / Glasser) | Influence Type | Extent (mm) | Magnitude |
| --- | --- | --- | --- | --- |
| 23949 | VisCent / V1 | Facilitative | 60.4 | 1.00 |
| 23907 | VisCent / V1 | Suppressive | 20.1 | 0.83 |
| 26109 | DefaultC / POS1 | Suppressive | 50.0 | 1.00 |
| 14572 | DorsAttnA / LIPv | Facilitative | 56.9 | 0.62 |
| 6804 | SomMotA / Area 2 | Suppressive | 50.0 | 0.49 |
| 16123 | ContB / PGs | Suppressive | 35.3 | 0.40 |
| 25595 | VisPeri / V1 | Suppressive | 30.9 | 1.00 |
| 25005 | VisPeri / V2 | Suppressive | 14.8 | 1.00 |

The decomposition achieved a reconstruction that closely resembled the original retinotopic selectivity map (Figure 3B,D), preserving its key spatial features. The stepwise reconstruction further revealed how individual spatial components progressively shape the distribution of selectivity across the cortical surface, highlighting interpretable sources of local enhancement or suppression (Figure 3A). Quantitatively, the model explained 68.4% of the variance in the left hemisphere and 65.2% in the right hemisphere across individuals in the test set (HCP dataset [16]). This corresponds to 92.3% of the explainable variance, when compared to the ceiling established by the group-average map in the training set: 74.1% for the left hemisphere and 70.6% for the right. Notably, this performance was achieved using only 8 spatial components per hemisphere, corresponding to 24 free parameters: each component being defined by its location on the cortical surface, spatial extent of influence, and amplitude of modulation. It was not by design that each hemisphere ended up with the same number of components. This compression shows that much of the functional topography of retinotopic selectivity can be captured by a small number of robust spatially organized influences.

**Figure 3.**
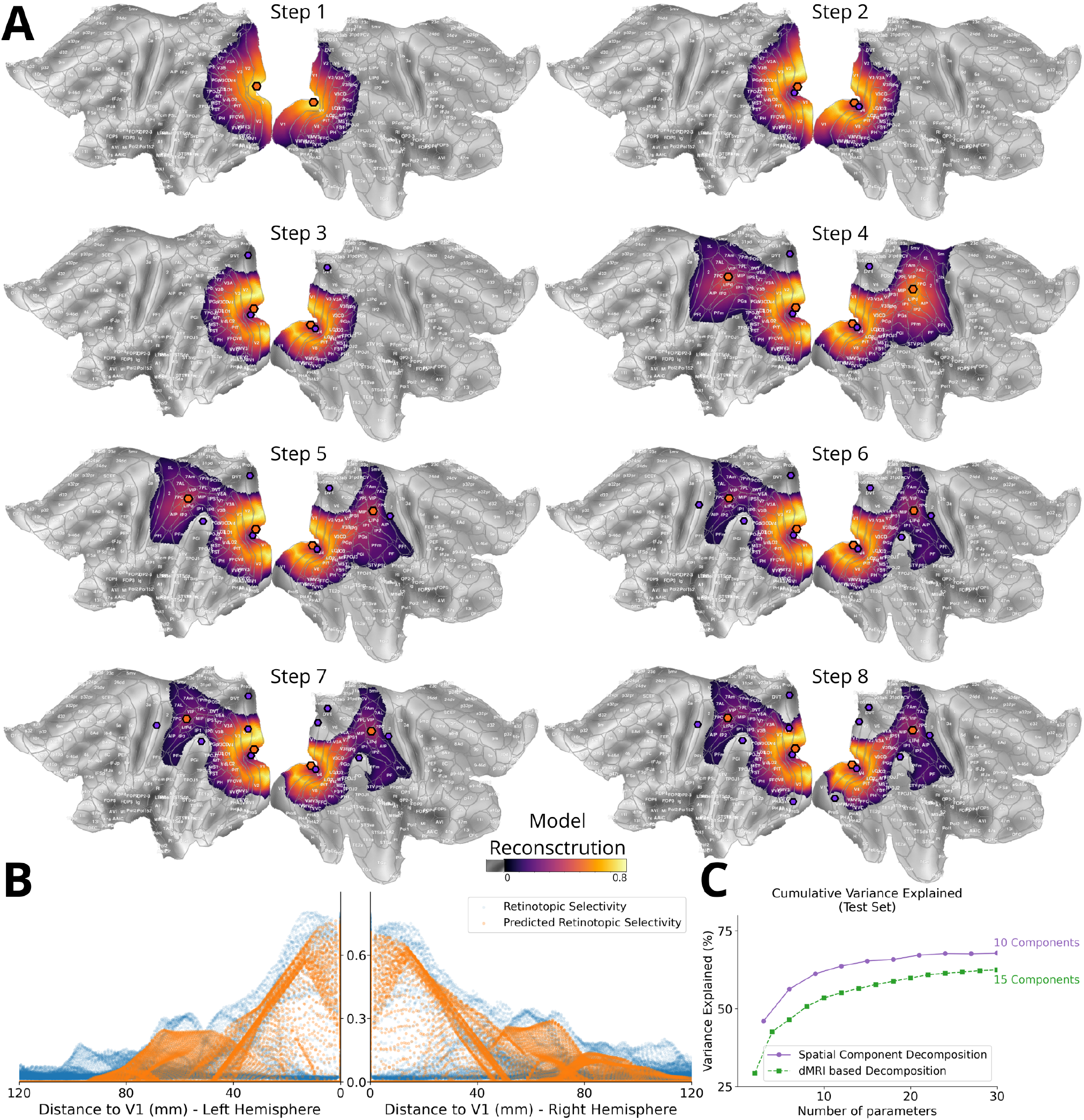
Stepwise Reconstruction of Retinotopic Selectivity on the Test Set. **A**. Incremental reconstruction of the test set for both hemispheres using spatial components selected during training. Components are added sequentially, with their cortical sources displayed as they contribute to the reconstruction. Regions with zero predicted selectivity are overlaid with sulcal patterns for anatomical reference. **B**. Scatter plots comparing the final model predictions to the actual retinotopic selectivity across cortical vertices, as well as to geodesic distance from V1, separately for each hemisphere. **C**. Cumulative variance explained on the test set as a function of model complexity (number of free parameters). The **purple** line corresponds to the Spatial Component Decomposition (SCD), where each component contributes three parameters; the **green** dashed line shows a dMRI tractography-based sparse dictionary model, with two parameters per component. Dots indicate successive components; final points (annotated) represent 10 SCD components and 15 dMRI-based components, respectively.

We compared SCD against a matched sparse dictionary derived from diffusion MRI-based structural connectivity [33]. Using the same number of parameters, this model explained 63.9% of the variance (Figure 3C), underscoring the predictive power of local geometry over long-range connectivity. The structure of the learned atoms and the resulting model reconstruction are shown in Supplementary Figure 3. Still, future extensions may combine SCD with sparse anatomical links [34] to further improve performance. Having established the accuracy of the SCD model, we next examined the anatomical and functional profiles of the identified spatial components.

### Competing Influences Within V1

To better understand these spatial influences, we now present and discuss the individual identified components, starting with those arising in primary visual cortex (V1). Given its dense thalamic input from the lateral geniculate nucleus (LGN)[24], V1 is a natural candidate for exerting strong spatial influences on retinotopic selectivity. Indeed, a third of the identified components originate within the anatomical boundaries of V1, including a dominant facilitative source (Figure 4A) and two pronounced suppressive ones (Figure 4B) in each hemisphere.

**Figure 4.**
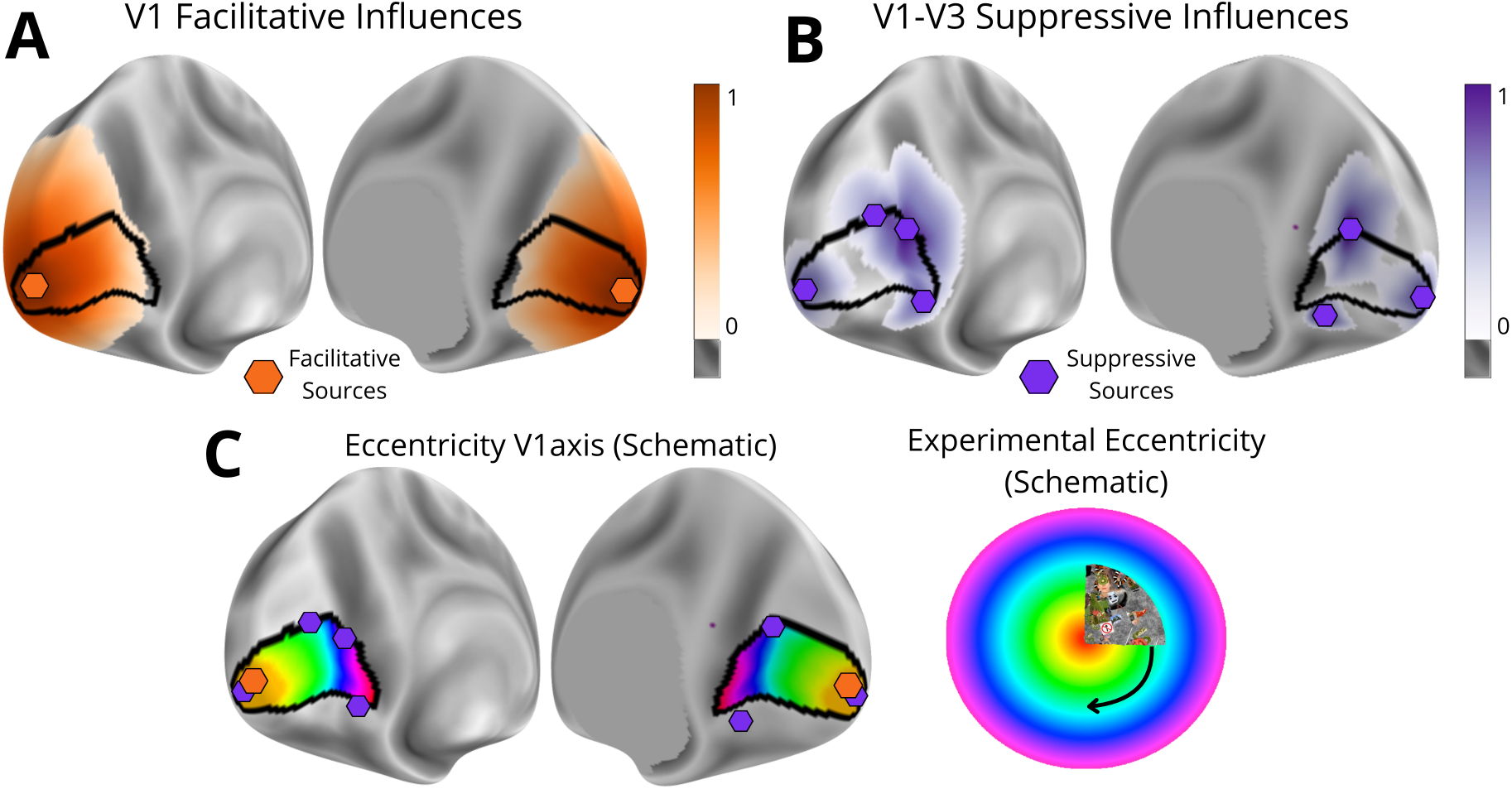
V1 antero-posterior eccentricity axis. **A** V1 facilitative influences displayed on the inflated cortical surface, with the V1 border overlaid for reference. **B** V1-V3 suppressive influences displayed on the inflated cortical surface, with the V1 border overlaid for reference. **C** Conceptual schematic representation of the antero-posterior eccentricity axis in primary visual cortex (V1), overlaid with the locations of all identified suppressive components. Right. Illustration of the visual stimuli used in the experimental paradigm (adapted from Benson et al., 2018), highlighting the limited eccentricity range (16°), which fails to drive responses in peripheral V1.

In each Hemisphere, the first of these suppressive components aligns anatomically with peripheral eccentricity zones along the anteroposterior axis of V1 [42, 43], where foveal regions reside at the occipital pole and peripheral fields map anteriorly (Figure 4C). Although human vision spans up to 60° of eccentricity, the stimuli in our dataset covered only the central 16° [16] (Figure 4D). As expected, anterior V1 showed little selectivity, but unexpectedly, this region exerted a suppressive influence extending over 30 mm (Figure 4A). This suggests that even in the absence of sensory-driven input from the thalamus, peripheral regions actively constrain the spatial spread of foveal representations—likely reflecting prior activity that has reshaped the functional landscape and continues to influence responses during this experiment.

The second suppressive component originated from the occipital pole—a region associated with foveal vision and therefor not attributable to the peripheral suppression described above. [42, 43]. Despite its strong retinotopic selectivity, the occipital pole exhibited markedly lower average functional connectivity compared to the rest of the cortex. Yet, this connectivity was not random: it was selectively structured toward transmodal systems, including the Default Mode and Control networks (Figure 5A, left). To probe this functional heterogeneity, we performed unsupervised clustering of whole-cortex connectivity profiles across V1 vertices (Figure 5B,C), revealing a bipartite organization: a dominant sensory-connected cluster, and a focal cluster at the occipital pole with transmodal-like connectivity (Figure 5D). This latter region coincided precisely with the origin of the focal suppressive component, suggesting that foveal V1 may be subject to additional competitive influences from transmodal systems.

**Figure 5.**
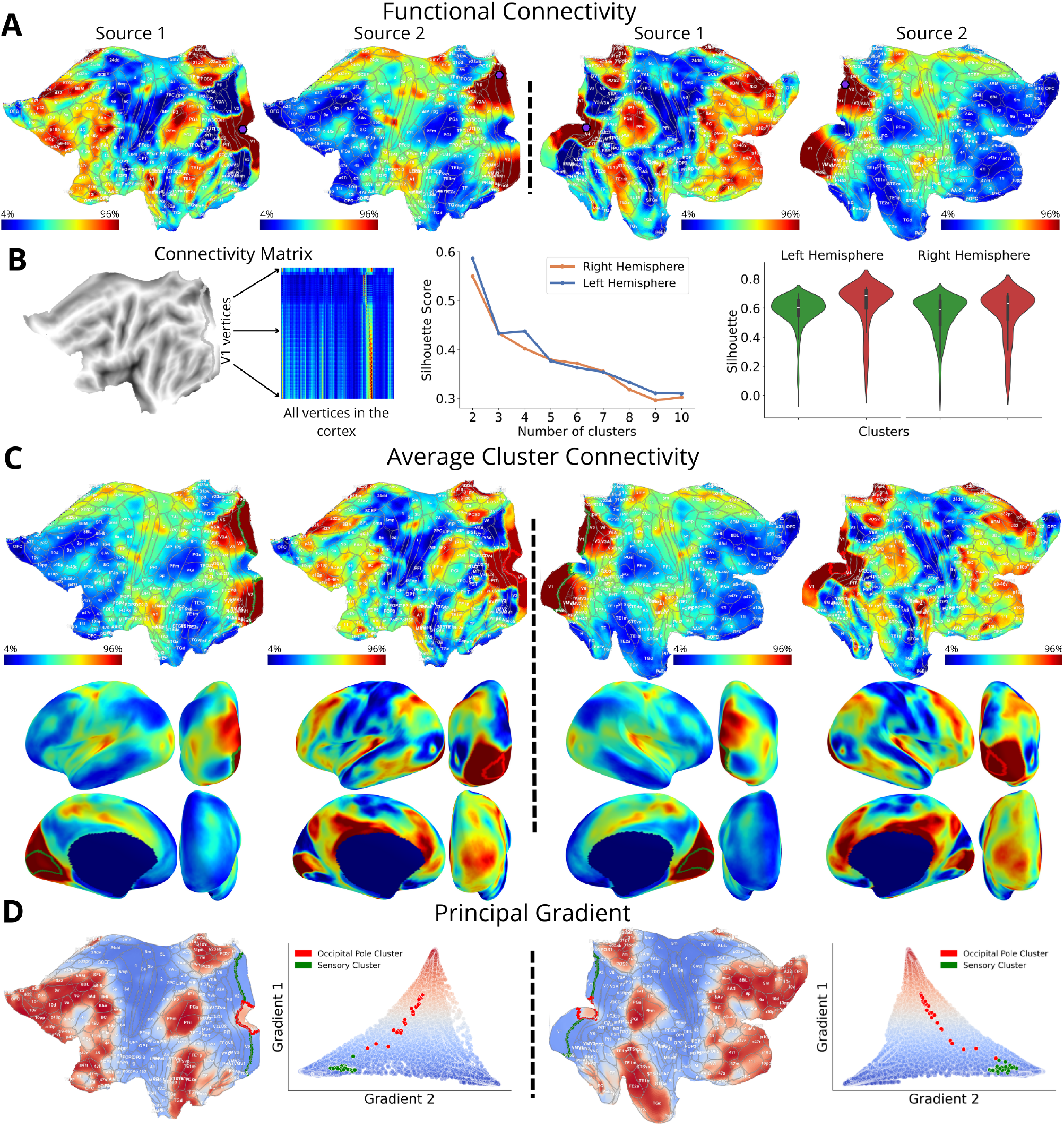
Two distinct functional clusters within V1. **A** Functional connectivity maps for each of the two V1-based suppressive sources, showing distinct patterns of functional connectivity. **B** (left panel) Functional connectivity matrix between all V1 vertices (rows) and the rest of the cortex (columns), sorted by cluster assignment. (middle panel) Silhouette score as a function of the number of clusters used in k-means clustering. (right panel) Distribution of silhouette scores within the two selected clusters (bottom). **C** Average whole-brain functional connectivity profiles for each V1 cluster. **D** Principal functional connectivity gradient (G1) from Margulies et al. (2016), with the two V1 clusters overlaid in red and green.

Projecting all V1 vertices into the space of principal functional connectivity gradients [44], the occipital-pole cluster aligned with a localized transmodal peak (Figure 5), reinforcing its deviation from classical unimodal organization. Previous studies have demonstrated eccentricity-dependent differences in V1 connectivity[45, 46], laying the groundwork for understanding functional heterogeneity within V1. Building on this foundation, our results reveal a focal foveal-pole subzone with a distinct transmodal fingerprint, pointing to a potential interface between retinotopic coding and higher-order systems at the core of visual input. Remarkably, this region may be analogous to higher-cognitive subzones recently identified in primary motor cortex [47], suggesting a shared organizational motif within unimodal systems.

### Suppressive Influences Arise from Primary Sensory Regions and Default Mode Network

In the previous section, we observed that the two dominant suppressive sources influencing V1 had very different connectivity profiles: one located in a primary sensory region, the other in a region strongly connected to transmodal hubs[48]. To examine their topological organization, we projected all suppressive components into the two-dimensional functional connectivity embedding defined by the first two principal gradients of cortical organization [44]. All sources clustered near the outer apex of this manifold (Figure 6A), corresponding to the functional periphery of cortex. The three outer peaks of this manifold are known to house 1) the Visual network 2) the Somatomotor-Auditory network and 3) the Default Mode Network [44]. A randomization test confirmed that the observed clustering of suppressive sources near the outer apex of the functional connectivity manifold—spanning all three peripheral peaks associated with the Visual, Somatomotor-Auditory, and Default Mode networks—was unlikely under chance (*p* = 0.037), indicating a significant bias toward regions at the extremes of functional differentiation.

**Figure 6.**
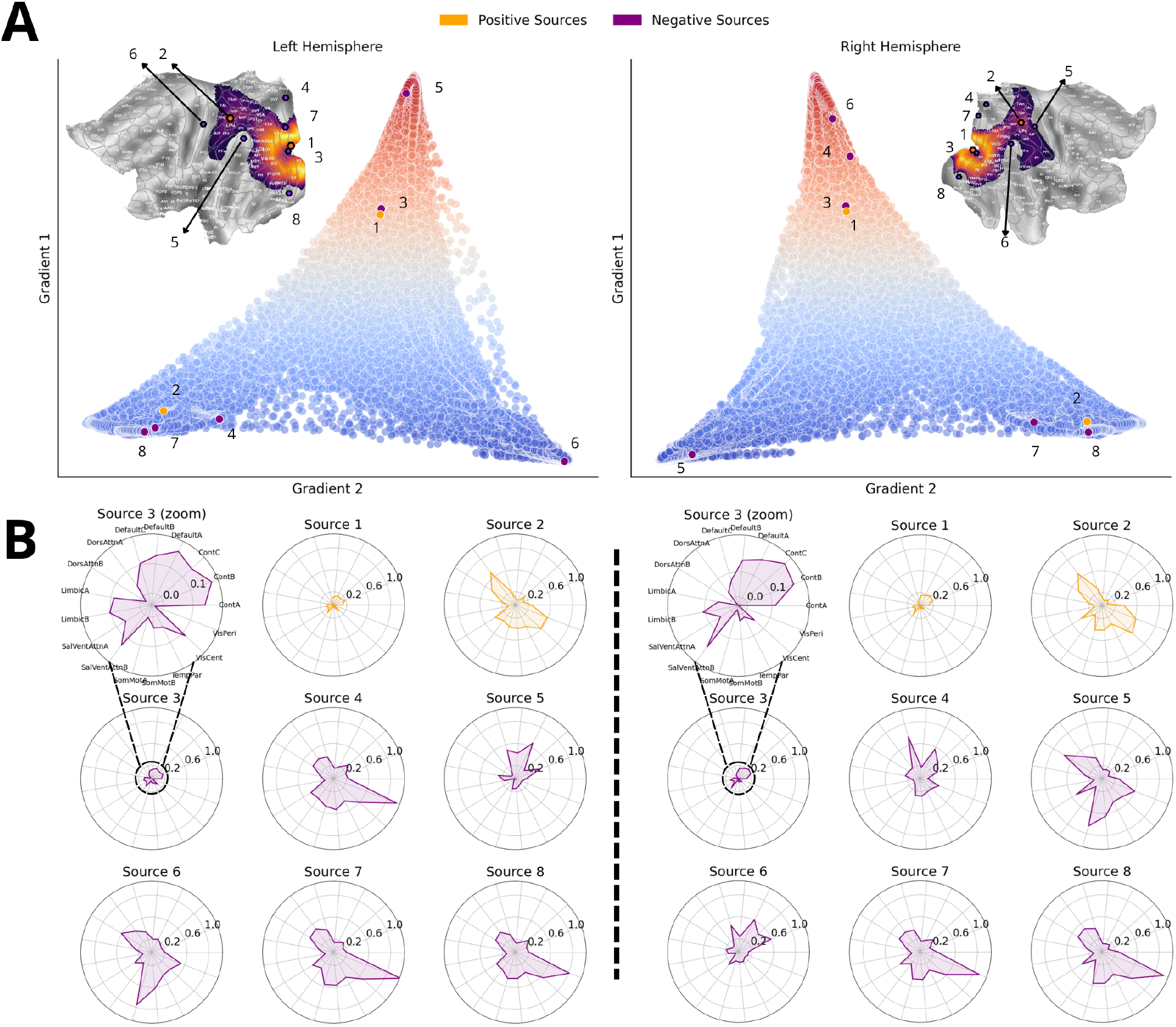
Suppressive Influences Arise from Primary Sensory Regions and the Default Mode Network. **A**. Locations of facilitative (orange) and suppressive (purple) influence sources projected into the 2D functional connectivity embedding space derived from Margulies et al. (2016) and on the flat map. **B**. Median functional connectivity profiles for each spatial source, visualized as spider plots across the 17 canonical networks defined by the Yeo atlas. These profiles highlight selective coupling to primary sensory and default mode systems. Source 3 shown in an enlarged inset to better illustrate its functional connectivity profile.

We wanted to test whether this pattern reflects a more general organizational principle. To this end, we examined the functional connectivity profiles of each suppressive component relative to the Yeo 17-network parcellation[49] (Figure 6B). Strikingly, every suppressive influence was strongly coupled either to competing primary sensory systems—such as the somatomotor network or peripheral visual regions—or to transmodal hubs, particularly within the default mode network (DMN). A permutation-based statistical test confirmed that these sources were significantly more connected to these three networks (somatomotor, peripheral visual and DMN) than expected by chance (left hemisphere: *p <* 10^−3^; right hemisphere: *p* = .013; see Methods for details). These findings support the interpretation that suppressive components reflect spatial competition 1) between sensory representations and 2) between sensory and transmodal systems.

These results suggest that competition for cortical territory is not limited to within-area dynamics or homogeneous sensory zones, but is a pervasive process that acts across distinct representational systems. Suppressive influences emerge at the hearth of both unimodal and transmodal networks, indicating that cortical function is shaped by long-range spatial competition. Notably, the spatial extent of these suppressive components—particularly those in transmodal and somatomotor regions—ranges from 30 to 50 mm, extending well beyond the classical boundaries of these areas and their associated networks. Retinotopic selectivity, rather than being embedded solely within a visual hierarchy, is subject to constraints imposed by global patterns of representational competition. Such cross-domain interactions reflect a broader organizational principle—wherein cortical specialization emerges not only from feedforward inputs or local gradients, but from the negotiation of influence among large-scale cortical systems.

## Discussion

### Retinotopic Information Spreads Across Cortex Through Spatial Influences

A central challenge in systems neuroscience is to understand how complex, high-dimensional representations are organized across the two-dimensional cortical sheet. While the local structure of sensory maps such as retinotopy has been extensively characterized, the global principles that govern their spatial distribution across the cortical surface remain poorly understood. Prior models have largely focused on discrete cortical areas or emphasized hierarchical connectivity[50], with limited attention to how representational strength varies spatially beyond early sensory regions. Even studies that have highlighted the widespread nature of retinotopic representations[17, 18] have typically treated these maps as isolated units rather than as components of a continuous topographic system.

In this study, we addressed this gap by introducing a quantitative framework for analyzing the macroscale organization of retinotopic selectivity across the human cortex. Using data from the Human Connectome Project, we first demonstrated that retinotopic selectivity—defined as the proportion of signal variance explained by a region’s preferred visual field location—declines gradually with geodesic distance from primary visual cortex (V1), forming a continuous spatial gradient. This result supports the idea that visuospatial dependence is not confined to isolated visual maps but extends across a broad swath of cortex in a spatially graded manner.

To further capture deviations from this monotonic decay, we developed Spatial Component Decomposition (SCD), a novel analytic method that models cortical maps as the superposition of spatially localized influences. Applied to retinotopic selectivity, SCD revealed a small number of spatial components—some additive, others suppressive—that together explained over 92.3% of the explainable variance in the data. Additive components emerged from early visual areas and parietal regions, while suppressive components are located in competing primary sensory regions and transmodal hubs[48], including nodes of the default mode network. These findings suggest that retinotopic encoding is not merely supported by sensory drive but is shaped by a competitive spatial landscape across functionally heterogeneous systems.

Notably, the original dataset contained about 30,000 cortical vertices per hemisphere-each with its own retinotopic selectivity value—yet the spatial component decomposition reduced this complexity to just 24 interpretable parameters per hemisphere. This finding suggests that the high-dimensional organization of cortical function may emerge from a relatively compact set of spatial influences. Such simplicity is consistent with self-organizing principles in which large-scale cortical patterns arise either from smooth, spatially structured gradients—potentially rooted in early developmental morphogen fields—or from the propagation of activity from focal cortical sources that gradually reshape the surrounding functional architecture.

Other authors have emphasised the spatial relationships between positive and negative activations across a range of tasks[51, 52, 53]. Notably, Leech et al. (2023)[53] identified that the spatial influences on functional connectivity differed across the cortex. This may be related to the different spatial extents in representations that we see in this study. It has also been shown, using a method adapted from earth sciences known as kriging, that positive activations are predictable based on the spatial layout of negative activations due to regularities in the spatial pattern across the cortex (Leech et al. (2024)[52]). Future work could apply kriging to test whether the patterns of suppressive and facilitative sources in this work are predictable based on spatial regularities (rather than cognitive networks). However, we also point out some important distinctions between the above and the present work. Here we focus on functional specialisation, a non-negative quantity, rather than positive and negative activations. The suppressive sources in this work do not represent negative activations (or negative representations), but chip away at the isotropic spatial influence of facilitative sources. Notably several of the suppressive sources are in areas that are positively activated by visual stimuli and have significant retinotopy (e.g. in V1), that is however lower than expected. Therefore suppressive sources and task-negative activity should not be confused. The finding of suppressive spatial influences of Default Mode nodes on visual retinotopy is unexpected and supports a competitive view of functional brain organization.

### Competition for Cortical Territory

Our findings support a model in which cortical representations emerge not solely from local feedforward inputs but through dynamic interactions among multiple functional systems, each exerting spatial influence over its surrounding cortical sheet. In this view, the formation of a mental representation is initiated by patterned input from subcortical sources—such as thalamic projections conveying sensory signals or hippocampal inputs contributing mnemonic content—each with its own characteristic topological covariance structure (for example, in retinotopy, nearby receptive fields tend to be co-activated). These inputs provide a scaffold upon which cortical circuits refine their connectivity, shaped by Hebbian plasticity mechanisms that strengthen co-active pathways and encode the correlation structure of the input space. Such dynamics are consistent with computational models showing that, when spatial embedding and resource constraints are taken into account, networks self-organize into modular, mixed-selectivity architectures that resemble those observed in the primate brain [54].

Once established, different sources of information—sensory, mnemonic, cognitive—compete to occupy and stabilize cortical territory. This competition is not arbitrary but governed by both anatomical constraints and functional demands, resulting in a landscape of mutually reinforcing and suppressive influences. Our spatial component decomposition makes this competitive structure explicit: regions that contribute to retinotopic selectivity are flanked by zones of suppression, explaining the anisotropic spread of retinotopic selectivity from primary visual cortex. The systematic presence of suppressive components within the default mode network (DMN) suggests that even abstract, internally generated content can exert spatial pressure on cortical real estate, limiting the spread of stimulus-driven representations.

This framework echoes and extends earlier theoretical models such as the optimized four-stream model proposed by Aflalo and Graziano [55], in which discrete visual areas emerge from the spatial competition between multiple hierarchically organized representational streams. By assuming only the existence of high-level functional anchors and enforcing spatial continuity, their model showed that the known layout of macaque extrastriate areas could emerge without hard-coded boundaries—suggesting that visual areas themselves may arise as natural borderlands in a cortical competition for functional specialization. Similarly, our results show that competitive interactions between representational domains—here revealed through suppressive gradients—can account for the segmentation of cortical space into distinct functional territories. This suggests that the borders between cortical areas may not be static anatomical partitions, but emergent equilibria shaped by overlapping demands on the cortical sheet.

This perspective aligns with recent theoretical accounts positing that cortical organization reflects a negotiated equilibrium among multiple representational domains[56, 57]. While our study focuses on a canonical form of sensory representation—retinotopic encoding—the principles uncovered here are likely to generalize, analogous to how the brain contains other topographic maps with similar organizational properties, such as tonotopic maps in auditory cortex (in which neighboring neurons respond to similar sound frequencies) and somatotopic maps in somatosensory cortex (in which neighboring regions represent adjacent parts of the body). Future work should investigate how similar decomposition techniques might explain maps from other modalities (e.g., somatosensory, auditory), high-level visual properties (e.g., object shape, motion trajectories), or to abstract cognitive variables (e.g., memory traces, social inferences, imagined scenarios) that characterize transmodal zones like the DMN. Such investigations would offer a more comprehensive account of how representational pressures shape the functional topography of the brain, revealing general rules by which neural systems carve out and defend their spatial niches.

### Functional heterogeneity within V1

By clustering all V1 vertices according to their whole-brain functional connectivity, we identified two clusters with strikingly different connectivity profiles. The larger cluster encompassed most of V1 and was strongly coupled to visual regions, with weaker but positive connections to other sensory systems and attentional networks. In contrast, a focal cluster at the tip of the occipital pole showed the opposite pattern: its strongest connections were with transmodal hubs [48], including both the frontoparietal control network and the default mode network (DMN).

These results converge with previous studies that examined eccentricity-dependent functional connectivity in V1 [46, 45]. Both studies elegantly demonstrated heterogeneity along the antero–posterior axis of V1, but relied on manual subdivisions of eccentricity bands, in contrast to our agnostic, data-driven clustering. The largest of these studies, by Sims and colleagues, likewise identified two broad subdivisions of V1 and compared their connectivity to three canonical Yeo networks (DMN, control, salience). They reported that the control network was preferentially connected to posterior (central) V1, consistent with our finding that the occipital pole has a transmodal connectivity profile. At the same time, they observed that different subdivisions of the DMN connect preferentially with different eccentricity segments of V1. At first glance, this may seem at odds with our results. However, their analyses focused on comparing relative connectivity between networks, whereas we examined the within-cluster connectivity fingerprint. The most important distinction in our data is that the occipital pole cluster is less connected to the rest of the visual system than to transmodal hubs—an effect not directly addressed in previous comparisons.

Sims and colleagues also explored potential long-range white-matter pathways linking V1 to frontoparietal and DMN regions, raising candidate structural sources for the functional influences we identify here. A natural next step will be to systematically apply their tractographic methodology to the suppressive influences uncovered by SCD, in order to constrain the anatomical substrates of these long-range interactions.

These findings highlight that even within primary visual cortex, a focal subzone exhibits a transmodal fingerprint, pointing to an overlooked interface between retinotopic coding and higher-order systems. A parallel has recently been described in primary motor cortex [47], raising the exciting possibility that other early sensory cortices, such as A1, may also harbor focal transmodal subzones.

### Functional Gradients and Delineated Areas

A longstanding debate in systems neuroscience concerns whether the cortex is best understood as a mosaic of discrete areas or as a continuous surface shaped by gradients of functional specialization [58, 44, 27]. Our results, while motivated by a gradient-based perspective, do not require a strict opposition to areal models. On the contrary, the spatial component decomposition we introduce reveals a pattern of influence that both aligns with known functional areas—such as V1—and extends well beyond their traditionally defined anatomical boundaries. This suggests that functional specialization may propagate through continuous spatial gradients, even as local areal identities remain intact.

Strong evidence supports the existence of discrete cortical areas, defined by sharp transitions in cytoarchitecture, connectivity, or function[24]. Yet, these boundaries need not contradict the presence of functional gradients. Rather, the two may reflect different facets of cortical organization. Gradients might traverse multiple areas, integrating their contributions into coherent large-scale patterns. Alternatively, areas themselves may represent locally uniform zones within a broader gradient of representational tuning—as seen in the systematic decline of retinotopic selectivity from V1 through V2 and V3. The gradients revealed by our model, in this view, are not incompatible with arealization but may capture the supramodal structure that connects local modules into a unified functional networks.

Importantly, the spatial extent of the gradients we observe—often spanning more than 50 mm of cortical surface—rules out simple methodological artifacts as their source. The effects of fMRI spatial smoothing and group averaging are typically confined to scales of a few millimeters[41, 40]. The robustness and reach of the spatial influences identified by our model therefore point to genuine macroscale structure in cortical organization, not to artifacts of preprocessing. These findings underscore the value of modeling the cortex not as a mosaic of isolated modules, but as a continuous surface in which specialized zones emerge and interact within broader gradient-based patterns of organization.

## Limitations and Future Directions

While our findings offer a novel perspective on the spatial organization of retinotopic selectivity, several methodological and interpretive limitations should be acknowledged. First, our analysis relied on functional data projected onto an average cortical mesh. This approach facilitates group-level comparisons but may obscure individual variability in cortical folding and retinotopic organization. Future work could refine these estimates by applying spatial component decomposition directly to each subject’s native 2D cortical surface, potentially increasing spatial precision and capturing idiosyncratic anatomical influences.

Second, although the spatial influences identified by our decomposition model account for a large proportion of the variance in retinotopic selectivity, they represent correlational structure rather than direct causal effects. The presence of a spatial component in a particular region does not, by itself, establish that the region exerts a mechanistic influence on nearby retinotopic encoding. Establishing causal directionality will require perturbative methods—such as transcranial magnetic stimulation (TMS) or lesion studies—that can more directly test whether the identified regions modulate selectivity in surrounding cortex. In parallel, computational models could be developed to reproduce functional specialization maps from first principles, using mechanistically defined spatial influences to test whether the observed patterns can emerge from plausible developmental or activity-driven processes.

Finally, our broader interpretation—that cortical representations compete for territory across the cortical sheet—rests on a set of anatomical regularities observed in the distribution of suppressive influences. While these patterns are systematic and statistically robust, stronger support for this theory would require convergent evidence from other representational domains. Future studies should extend this framework to non-visual modalities, abstract cognitive content, or developmental trajectories to assess whether the competitive structuring we observe generalizes across functional systems. Such evidence would strengthen the claim that representational competition is a widespread organizing principle of cortical function.

## Conclusion

Our analysis reveals a detailed macroscale topographic account of how retinotopic selectivity is distributed across human cortex. A dominant facilitative influence originates in V1, extending over 60 mm across posterior cortex, consistent with its dense thalamic input and primary role in visual encoding. This widespread influence is sculpted by suppressive components arising both from competing primary sensory areas and from transmodal regions, particularly nodes of the default mode network (DMN). Despite the complexity of the cortical surface, these effects were captured with just 24 spatial parameters, explaining over 92.3% of the explainable variance. This suggests that retinotopic information is represented in a distributed and macroscale pattern shaped by competing cortical landmarks, revealing a spatial dimension to the well-established dynamic antagonism between sensory and transmodal systems[38, 59].

The method we introduce—Spatial Component Decomposition (SCD)—offers a flexible, interpretable framework for modeling cortical organization. By decomposing any functional map into spatially localized, signed influences, SCD provides a means to quantify how different regions contribute to, or suppress, a given feature. Unlike high-dimensional representations, SCD yields compact descriptions grounded in cortical geometry. Its generality makes it well-suited for analyzing other cortical features—from structural properties[60, 26, 12] to gradients of abstraction and self-related processing[61].

More broadly, our results suggest that cortical representations emerge through a form of spatial competition, where distinct informational domains fight for influence over cortical territory. This competition persists even in the absence of direct stimulus input: peripheral visual zones, despite lacking relevant stimuli, still constrain nearby representations; similarly, the DMN suppresses retinotopic encoding from the core of V1. Notably, this suppressive influence appears selective to the transmodal hubs[48], and more specifically the DMN; other high-level systems, such as the attention networks, did not produce comparable suppressive components or exert significant spatial constraints on retinotopic encoding. Our findings support the view that the brain’s functional architecture reflects a dynamic contest for cortical territory—where distant systems impose competing claims on the same neural real estate[62, 63, 64, 65].

## Methods

### Population Receptive Fields (pRFs)

Here we analyzed existing population receptive field data across the human cortex from two openly available datasets [16, 22]. Briefly, population receptive fields (pRFs) are a computational model used to characterize the spatial selectivity of neural populations in the visual cortex. A pRF describes the region of the visual field to which a given voxel or neural population is responsive, essentially summarizing how visual stimuli are represented in the brain. This is achieved by estimating key parameters: the pRF center (its spatial location in the visual field), size (the spread or extent of the receptive field), and amplitude (the strength of the neural response). These estimates are obtained by modeling the relationship between visual stimuli presented to participants and the recorded neural activity, typically using functional magnetic resonance imaging (fMRI) [13, 7]. By systematically presenting stimuli that sample the visual field, pRF models provide voxel-wise information about visual field representations, offering insights into the functional architecture and retinotopic organization of the whole cortex.

#### Data Availability

In this study, we used preprocessed pRF datasets from publicly available neuroimaging repositories, specifically the Human Connectome Project (HCP) retinotopic dataset [16] and the NYU retinotopic dataset [22]. These datasets provide precomputed pRF parameters for individual participants, which were derived from standard pRF modeling algorithms applied to high-resolution functional magnetic resonance imaging (fMRI) data. The datasets used in this study are publicly available and can be accessed from their respective repositories. The Human Connectome Project (HCP) dataset can be downloaded from https://db.humanconnectome.org/, the NYU dataset is available at https://osf.io/bw9ec/.

#### pRF Parameters

Population receptive field (pRF) models provide voxel-wise estimates of:

- **Receptive field center** (*x*_0_, *y*_0_): Cartesian coordinates in visual field space specifying the spatial location of each voxel’s receptive field.
- **Receptive field size** (*σ*): The standard deviation of the Gaussian profile, representing the spatial extent of the receptive field.
- **Receptive field magnitude**: A measure of the amplitude or strength of the response within the receptive field, reflecting the voxel’s sensitivity to the stimulus at its preferred location.
- **Variance explained**: A goodness-of-fit measure indicating the proportion of signal variance in the fMRI BOLD-activity time course that is accounted for by the pRF model at every vertex.

These parameters are obtained using the standard pRF modeling approach described by Dumoulin and Wandell [13], incorporating the convolution of predicted neural responses with the canonical hemodynamic response function (HRF) (Supplementary Figure 1).

#### HCP dataset experimental procedure

##### Experimental Procedure

The primary dataset used in this study is the HCP retinotopy dataset [16]. This dataset includes pRF maps extracted from high-resolution 7T fMRI data. The study included 181 subjects, scanned using a 7T fMRI scanner with a 1.6 mm isotropic voxel size and a TR of 1s. Participants viewed dynamic visual stimuli consisting of moving apertures with a maximum eccentricity of 8 degrees. Each participant completed six 5-minute runs, totaling 30 minutes of recording.

##### Preprocessing

The preprocessing of this dataset followed the HCP pipeline, which projects the data into the fs lr32 cortical surface space. Denoising was performed using multirun spatial Independent Component Analysis (sICA) combined with FIX. The pRF extraction used an isotropic Gaussian pRF model within the fs lr32 cortical space [66].

#### Additional Data Preparation

To prepare the retinotopic selectivity data for analysis, we implemented a custom pipeline that processes data from the original pRF model outputs into normalized and partitioned datasets (Supplementary Figure 1). First, we excluded the medial wall, retaining only cortical vertices. Min-max normalization was performed for each subject individually to preserve within-subject variability in retinotopic selectivity values while ensuring consistent scaling across subjects.

#### Retinotopic Selectivity

Retinotopic selectivity quantifies the degree to which neural activity at each cortical vertex is driven by retinotopic information. Specifically, it measures the fraction of the total variance in the recorded signal that can be attributed to the predicted responses based on visual field mapping. High retinotopic selectivity indicates that a vertex is strongly driven by spatially localized stimuli within its receptive field, reflecting precise retinotopic organization. This metric is typically expressed as the variance explained (*R*^2^) at each vertex, with higher values indicating a robust match between the observed neural responses and the modeled retinotopic representation. It is important to note that retinotopic selectivity does not correspond to cortical activity but rather to functional specialization; a region can exhibit high retinotopic selectivity with a small fMRI signal if its activity is primarily driven by retinotopic information but remains low compared to baseline activity.

#### Train, Test, Validation

To facilitate model training and evaluation, we partitioned the HCP dataset into separate training and test subsets, using an 80/20 split ratio across participants. This approach allowed us to assess generalization performance within the same acquisition domain. Additionally, we used the independent NYU retinotopic dataset as a validation set, leveraging its distinct acquisition characteristics to identify and retain only the most robust and generalizable spatial components (atoms). This cross-dataset validation helped mitigate overfitting and ensured that our decomposition captured spatial structures that were consistent across different retinotopic mapping protocols.

#### Ceiling Computation for Visualization

To improve contrast in the visualization of the retinotopic selectivity map (Figure 1B), we applied a ceiling based on the empirical distribution of variance-explained values across vertices, following the same procedure reported in the original study [16]. We fit a two-component Gaussian Mixture Model (GMM) using GaussianMixture from the sklearn package [67]. The model identified one component with a lower mean (interpreted as noise) and one with a higher mean (interpreted as signal). We then computed posterior probabilities over a fine grid and selected the crossover point—where the posterior probability of the higher-mean component exceeds that of the lower—as the threshold. This yielded a ceiling of 9.8% variance explained, applied only for visualization. All model fitting and statistical analyses were conducted on the full, unthresholded data.

### Geodesic Distance

Geodesic distance refers to the shortest path between two points along a curved surface, such as the brain’s cortical surface, which is highly folded and convoluted. Unlike straight-line Euclidean distance, geodesic distance respects the intrinsic geometry of the cortex, making it a more accurate measure of spatial relationships and potentially reflecting the propagation of neural activity through local grey matter connections rather than long-range white matter tracts. Geodesic distances are computed within each hemisphere and do not cross the interhemispheric boundary. In this study, we used the surfdist Python package [68] to calculate these distances efficiently along cortical surface meshes.

### Spatial Component Decomposition

#### A Fourier-Inspired Approach for Cortical Map Decomposition

##### Spatial Component Decomposition

(SCD) is a novel method for decomposing high-dimensional cortical maps into a set of spatially localized components. SCD is derived from sparse dictionary learning principles but is specifically adapted to the irregular geometry of the cortical surface. Each component, or atom, is modeled as a threshold-linear function of the geodesic distance from a vertex, ensuring spatial localization while allowing for flexible, data-driven representations. Unlike traditional Fourier decompositions, which impose orthogonality and global basis functions, SCD learns a sparse set of spatial components that can independently capture facilitative or suppressive influences in the data. Applying SCD to functional specialization maps enables a compact and interpretable representation of their spatial structure. Critically, SCD can reveal both reinforcing and antagonistic spatial influences that may be obscured in traditional statistical maps, offering novel insights into the organization of cortical functional specialization.

#### Input Maps for Spatial Component Decomposition

The input data for **SCD** consists of statistical maps derived from neuroimaging analyses, representing measures of functional specialization across the cortical surface. Each map provides a value at every vertex of the cortical surface, with a resolution of 32,492 vertices per hemisphere using the chosen mesh. These maps are collected across N subjects, capturing inter-subject variability while maintaining a consistent spatial structure for analysis.

#### Sparse dictionary learning (SDL)

SDL aims to represent data as sparse combinations of learned atoms, providing flexible and compact representations. Unlike fixed bases such as Fourier transforms, SDL learns an overcomplete dictionary directly from the data, balancing reconstruction accuracy with sparsity. The underlying optimization problem involves minimizing reconstruction error while enforcing sparsity via the *l*_0_-norm, making the problem non-convex and computationally intractable in general. Consequently, practical solutions rely on heuristics to obtain approximate but efficient solutions. This approach enables improved interpretability and efficiency in tasks such as image compression, signal recovery, and feature extraction [69, 70, 71].

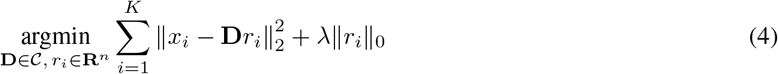

In this formula,

- *x*_*i*_ ∈ ℝ^*d*^ is the *i*^th^ data vector (here, a vertex-wise retinotopic-selectivity map),
- **D** ∈ ℝ^*d×n*^ is the dictionary matrix. Each column is a spatial atom (component),
- 𝒞 is the set of all admissible dictionaries (here, all possible combinations of component source on the cortical surface (anchor vertices) and spatial extents),
- *r*_*i*_ ∈ ℝ^*n*^ is a sparse coefficient vector that encodes *x*_*i*_ in the dictionary basis,
- *K* is the number of training samples (subjects),
- ∥ · ∥_2_ denotes the Euclidean norm,
- ∥ · ∥_0_ counts the number of non-zero entries (the *ℓ*_0_ “norm”), and
- *λ >* 0 is a hyper-parameter that balances reconstruction fidelity against sparsity.

In our adaption of SDL, we constrain the predictions to be positive using a ReLU function, ensuring that components can only suppress the underlying signal, mimicking the effect of local inhibitory influences on a positive baseline map.

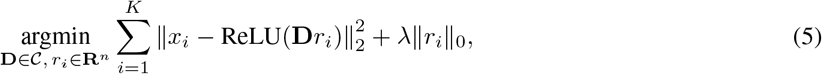

#### Atom definition

The atoms used in **Spatial Component Decomposition** are spatially localized functions centered on a set of vertices and defined on the cortical surface. In principle, these atoms can belong to any class of functions, such as Gaussian functions, decaying exponential functions, or any other form, provided they exhibit a spatially localized range. For this work, we specifically use threshold linear functions based on geodesic distance, which are computationally efficient and geometrically intuitive. Let **d**(*v, v*_0_) represent the geodesic distance between a cortical vertex *v* and a atom origin vertex *v*_0_. The atom *k*(*v*; *v*_0_) is defined as:

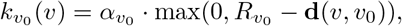

where 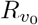 is the atom’s spatial range parameter, controlling the size of the localized region influenced by the atom, and 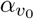 is a scaling factor that determines the slope of the atom’s influence. atoms with negative contributions are similarly defined as:

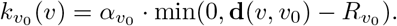

These threshold linear atoms ensure that each component is spatially localized, with the influence of the atom decaying linearly to zero at a distance of 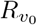. This choice of atom allows for the decomposition of statistical maps into spatially constrained components that are biologically interpretable and computationally efficient to calculate.

#### Loss Function

The effectiveness of the **Spatial Component Decomposition** method is evaluated through a loss function that quantifies the proportion of variance explained by the reconstructed maps relative to the original statistical maps. This is computed as a percentage of the variance captured by the reconstruction.

Let *M*_*s*_ represent the original statistical map for subject *s*, where *M*_*s*_(*v*) is the corresponding statistic at vertex *v*. Let 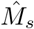 represent the reconstructed map obtained from the decomposition, where 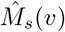 is the predicted statistic at the same vertex *v*. The coefficient of determination 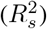 for subject *s* is defined as:

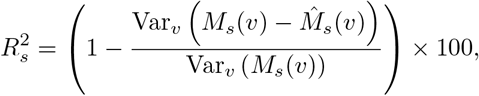

where Var_*v*_ (·) denotes the variance across all vertices *v* on the cortical surface.

The loss function is then defined as the average proportion of variance not explained across all subjects in the dataset (e.g., training, validation, or test set). To minimize the loss, we maximize the variance explained. Formally, the loss function is:

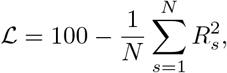

where *N* is the number of subjects.

#### Interpretation and Benefits

This loss function ensures that the reconstructed maps minimize the residual variance relative to the total variance of the original maps. By maximizing the percentage of variance explained, the method ensures that the spatial components accurately capture the essential patterns in the data. Evaluating ℒ across separate datasets (training, validation, and test) provides a clear metric for comparing different models and assessing the robustness and generalizability of the decomposition.

#### Algorithm for Deriving Components

Sparse dictionary learning is a non-convex optimization problem, for which no closed-form or globally optimal solution can be computed efficiently. Classical greedy approaches, such as Matching Pursuit (MP), build sparse representations by iteratively selecting atoms from a predefined dictionary to minimize residual error. However, MP cannot be directly applied in our setting due to the presence of a ReLU nonlinearity in the prediction function, which enforces non-negativity and introduces asymmetry into the solution space. To address this, we developed a novel greedy heuristic, inspired by MP but adapted to accommodate these constraints.

We make use of two independent datasets: the Human Connectome Project 7T retinotopy dataset [16], comprising 181 subjects, and the NYU-retinotopy dataset [22], comprising 44 subjects. The dictionary learning procedure is performed exclusively on the Benson dataset, which we divide into separate training and test subsets. Atoms are extracted from the training set and evaluated on the test set using a variance-explained threshold to enforce sparsity. The resulting model is then transferred to the Himmelberg dataset for validation: we fix atom locations and refit only their parameters. Atoms that fail to generalize—i.e., whose removal does not significantly impair performance on the validation set—are pruned. This two-stage procedure enables us to identify a sparse, interpretable dictionary of components that is statistically robust and replicable across datasets.

The detailed heuristic proceeds in the following stages:

1. **Initialization and Atom Addition** We begin with the HCP 7T retinotopy dataset (Benson et al., 2018) as our **training dataset**, which we further divide into separate **training and test subsets**. Starting with an empty dictionary, atoms are added sequentially. At each step, candidate atoms are defined by their cortical location (vertex). For each candidate, we jointly refit the amplitudes of all existing atoms along with the candidate, and evaluate the total variance explained on the **training set**. The candidate that yields the greatest improvement in explained variance is selected and permanently added to the model. Following each accepted addition, we locally optimize the locations and extents of all existing atoms. Each atom’s vertex is perturbed within its direct neighbors on the cortical mesh to find a new position that improves the model’s fit. This local refinement helps escape discretization artifacts and improves the alignment between atom locations and underlying signal structure.
2. **Stopping Criterion** The impact of the newly added atom is then evaluated on the **test set** without additional fitting: only atoms that improve the variance explained on the test set by at least 0.1% (corresponding to *λ* = 0.001) are retained. This ensures early stopping and guards against overfitting. This process continues iteratively until no candidate atom can increase the variance explained on the test set by more than the threshold (0.1%). At this point, the model is considered fit to the source dataset, with a sparse dictionary of spatial components.
3. **Cross-Dataset Pruning via Permutation Testing** To assess the generalizability of the derived atoms, we transfer the full dictionary to an independent dataset [22] that serves as our validation dataset. In this transfer we keep each atom’s center fixed, but we re-estimate its spatial extent and amplitude, allowing us to cross-validate whether the same underlying sources of influence are present—even if their spread or strength differs across samples. We then evaluate each atom’s necessity using a leave-one-atom-out permutation test. For each atom, we compare the performance of the full model to that of a reduced model obtained by omitting that atom. For both models, we compute the loss for every subject in the training set, yielding two labeled sets of loss values. We then randomly shuffle these labels to generate a null distribution for the difference in mean loss between the two models. Atoms are retained only if their exclusion leads to a statistically significant decrease in variance explained (*p <* 0.005), Bonferroni-corrected for multiple comparisons, thereby pruning non-generalizable components.
4. **Final Model** This final model offers a compact and interpretable representation of functional organization in visual cortex that is robust to overfitting and supported by replication.

##### Algorithm 1

Greedy Heuristic for Sparse Spatial Decomposition with Validation

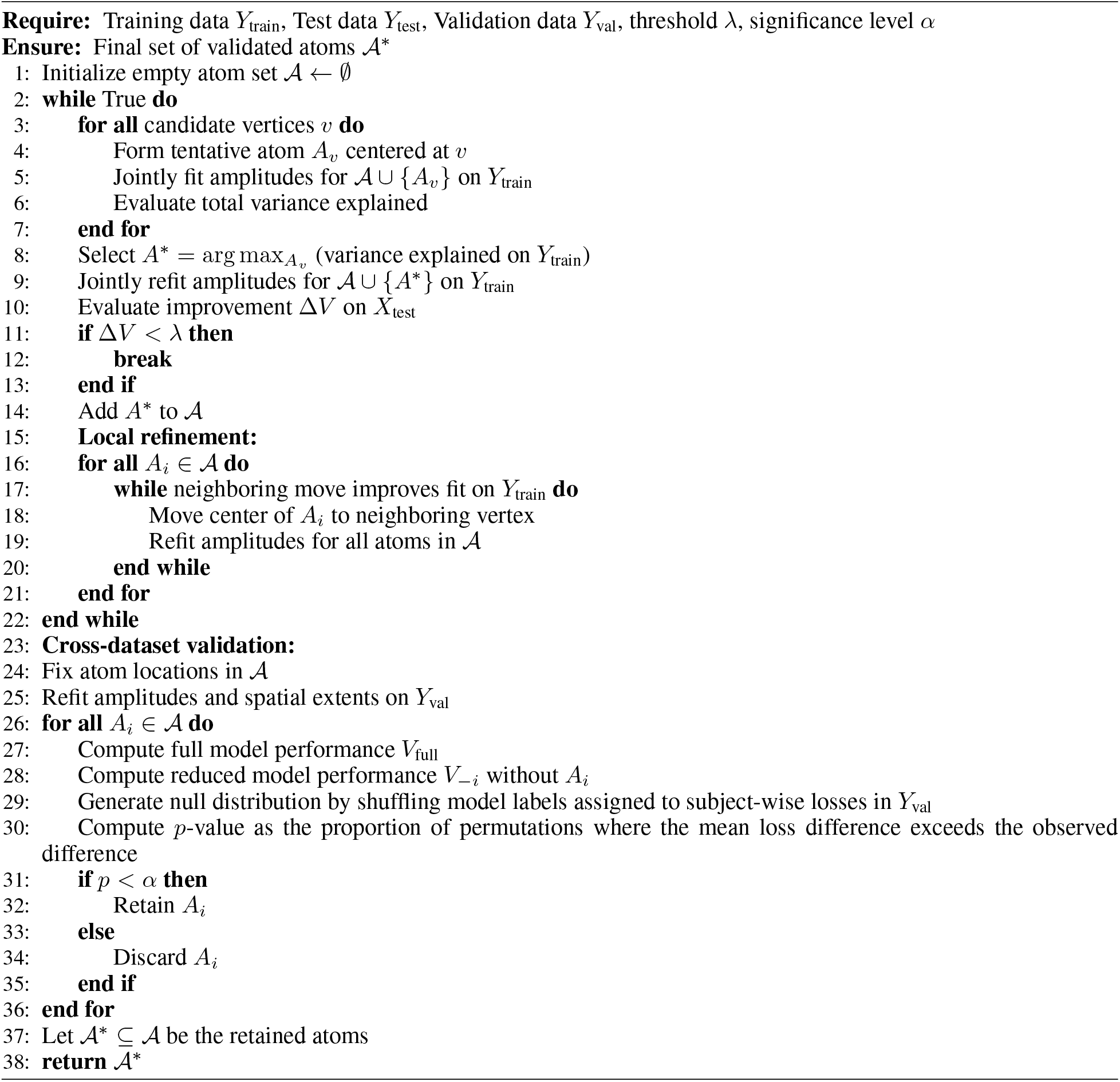

### Comparison with Tractography-Informed Decomposition

To compare our **Spatial Component Decomposition** (SCD) with anatomically-informed alternatives, we conducted an additional sparse dictionary learning analysis using structural connectivity profiles derived from tractography data. In this formulation, each atom corresponds to the structural connectivity profile of a single cortical vertex—that is, a vector indicating its probabilistic tractography-derived connectivity to all other vertices. We followed a similar greedy procedure as in SCD, sequentially adding atoms that best reduced the residual error in the training set.

Unlike the SCD model, however, no ReLU constraint was applied, and the atoms were directly derived from empirical connectivity data without any eigendecomposition or basis transformation. The model was trained on the same training subset of the HCP dataset, and performance was evaluated on the held-out test set using the same variance-explained metric described above. This structurally grounded decomposition allowed us to assess the extent to which the spatial organization of functional selectivity could be predicted by anatomical connectivity alone.

### Threshold-Linear Model

#### Model

For the naive threshold-linear model, we fit a three-parameter function with a maximum value, a slope, and a ceiling. This optimization problem is convex and was fitted using `jaxopt.ScipyMinimize` [72].

#### Loss function

The loss was defined as the mean variance explained across subjects:

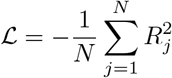

where 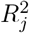 is the variance explained for subject *j*, and *N* is the number of subjects.

#### Residual Map

We computed a residual map by averaging, across subjects, the difference between the model prediction and the individual data.

### Principal Gradients

We used the first two cortical functional gradients computed by Margulies et al. (2016) on the *fs lr32* mesh. These correspond to the first two dimensions of a nonlinear dimensionality reduction performed with diffusion map embedding on resting-state fMRI data from the Human Connectome Project (HCP). These gradients are publicly available via neuromaps [73].

### Clustering of Functional Connectivity in V1

#### Functional Connectivity

To investigate the functional heterogeneity within primary visual cortex (V1), we analyzed whole-brain resting-state functional connectivity profiles for all V1 vertices. The functional connectivity data were derived from the Human Connectome Project (HCP), leveraging minimally preprocessed resting-state fMRI data.

#### Restriction to V1

V1 was defined using the **Glasser Atlas** [24] projected to the *f_lr32* mesh, yielding 831 vertices. We applied a PCA to reduce dimensionality, keeping 136 components combining 99.9% of the variance. We used `StandardScaler` and PCA from the `sklearn` package [67].

#### K-Means Clustering

We applied **K-means** clustering to the reduced dataset, testing *K* from 2 to 10. Cluster quality was assessed using the **silhouette score**:

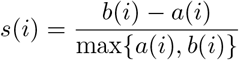

where *a*(*i*) is the mean intra-cluster distance, and *b*(*i*) the minimum mean distance to other clusters. The silhouette score quantifies how well each vertex fits within its cluster versus others. For the selected clustering at *K* = 2, we computed vertex-wise silhouette scores for both clusters.

### Null Model of Functional Network Assignment

To evaluate whether the observed suppressive components are unusually close, in functional connectivity space, to specific large-scale networks, we constructed a null model based on whole-brain connectivity profiles. For each of the six suppressive sources, we computed its median resting-state functional connectivity to each of the 17 canonical networks defined by the Yeo parcellation [49], yielding a coarse-grained connectivity profile across the cortex. Because our hypothesis focused on competition involving three systems—somatomotor, peripheral visual, and the default mode network (DMN)—we extracted the maximum functional connectivity of each source to these three networks. A high value on this measure indicates that a source is strongly connected to at least one of these networks, whereas a low value implies weak or diffuse connectivity across all three, suggesting little functional proximity to the hypothesized domains.

We then averaged this maximum connectivity value across the six suppressive sources, producing a summary measure of how strongly these components are functionally coupled to the targeted networks. To determine the statistical significance of this coupling, we generated a null distribution by repeatedly sampling six vertices at random from the cortical surface (1 million iterations), computing the same summary measure for each sample. We then calculated the proportion of random samples that yielded a higher average connectivity to the three networks than the observed value.

### Apex Proximity Statistical Test

To quantify how the suppressive components relate spatially to the boundary of the cortical functional embedding, we assessed both their proximity to, and coverage of, the embedding’s periphery. For each hemisphere, we first projected every cortical vertex into a 2D principal component space, where the xx-coordinate corresponds to its value on the first principal gradient and the yy-coordinate to its value on the second principal gradient (Figure 7B). The “anchors” of the embedding were defined as the three vertices lying at the outermost points of this manifold. We then considered only the suppressive sources (suppressive components) and computed two Euclidean distance metrics: (1) the maximum distance from each suppressive source to its nearest anchor, and (2) the maximum distance from each anchor to its nearest suppressive source. The first measures how close the suppressive sources lie to the cortical periphery; the second captures how well they span the full extent of the apex. To test whether this spatial configuration could occur by chance, we generated 10,000 random sets of six cortical vertices (matching the number of suppressive components) and computed the same two metrics for each set. We then calculated how often random configurations performed at least as well as the empirical one on both criteria.

**Figure 7.**
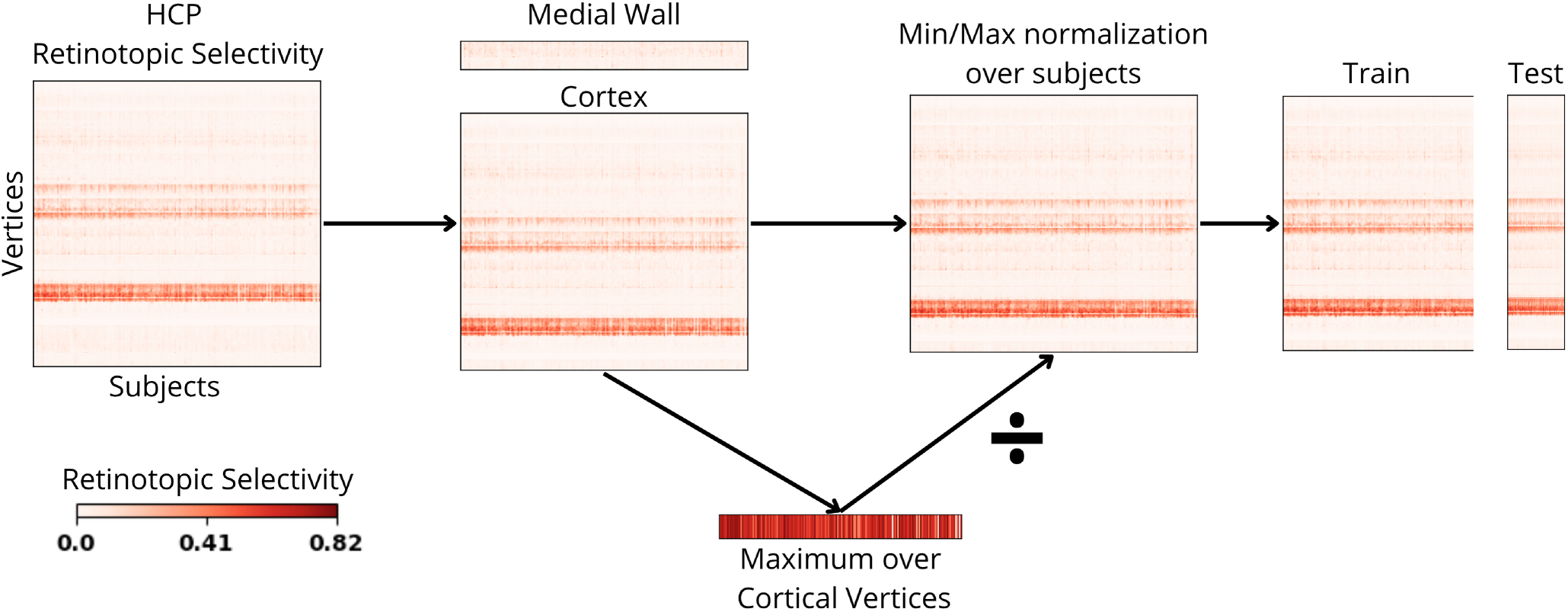
Supplementary Figure 1 Preprocessing pipeline for retinotopic selectivity data. Starting from the original pRF model outputs, we excluded the medial wall to retain only cortical vertices. Each subject’s data was then minmax normalized individually, preserving within-subject variability in retinotopic selectivity while enforcing consistent value scaling across subjects. The resulting datasets were then partitioned for subsequent analysis (see Supplementary Figure 1).

**Figure 8.**
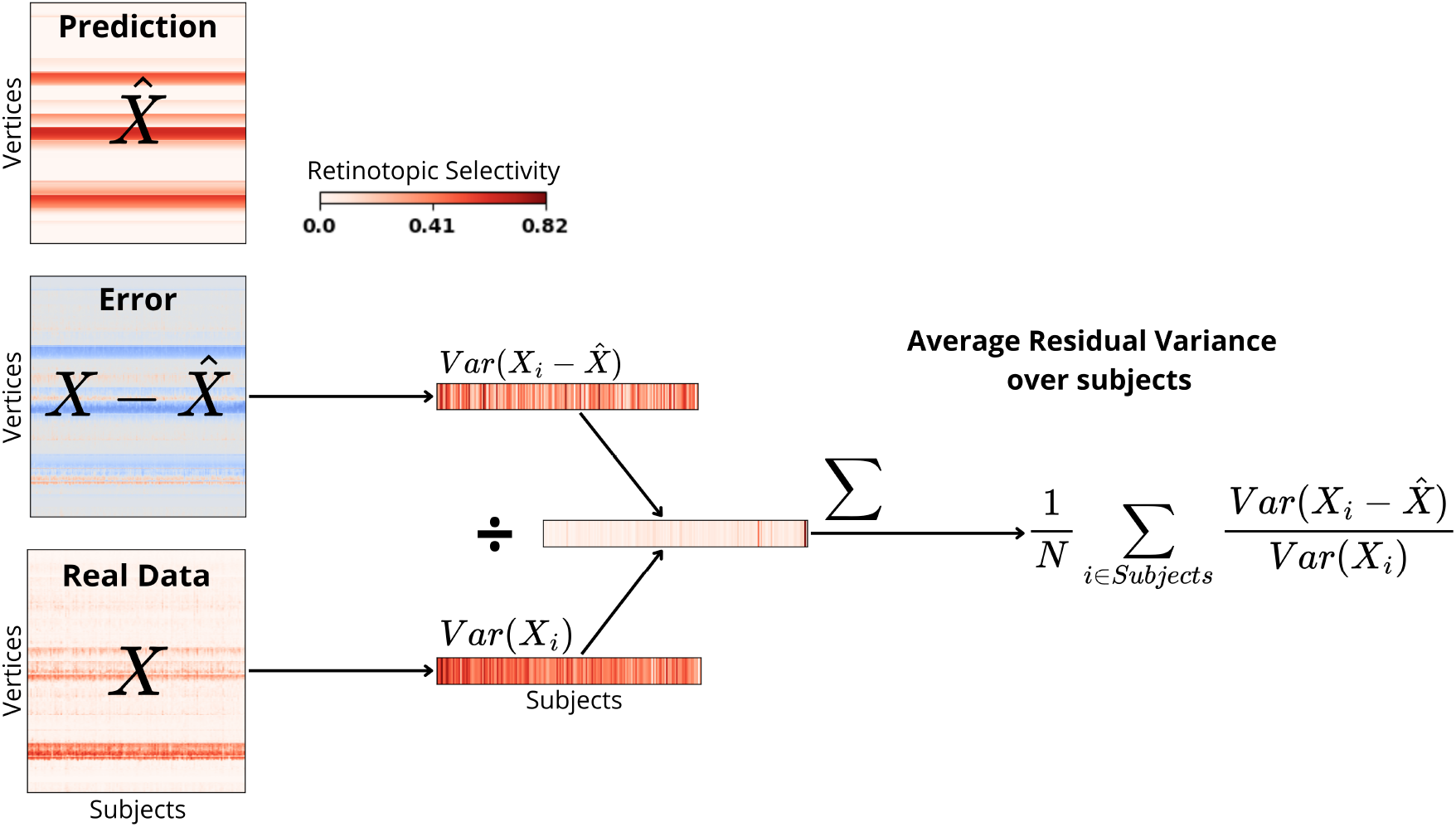
Supplementary Figure 2 Definition of the loss function used to evaluate model performance. The loss is computed as the average proportion of variance not explained across all subjects in a given dataset (training, validation, or test). For each subject, the variance explained (V E_s_) is defined as the proportion of variance in the original statistical map captured by the reconstructed map. Minimizing the loss function ℒ is equivalent to maximizing the average variance explained across subjects, ensuring accurate and generalizable spatial decompositions.

**Figure 9.**
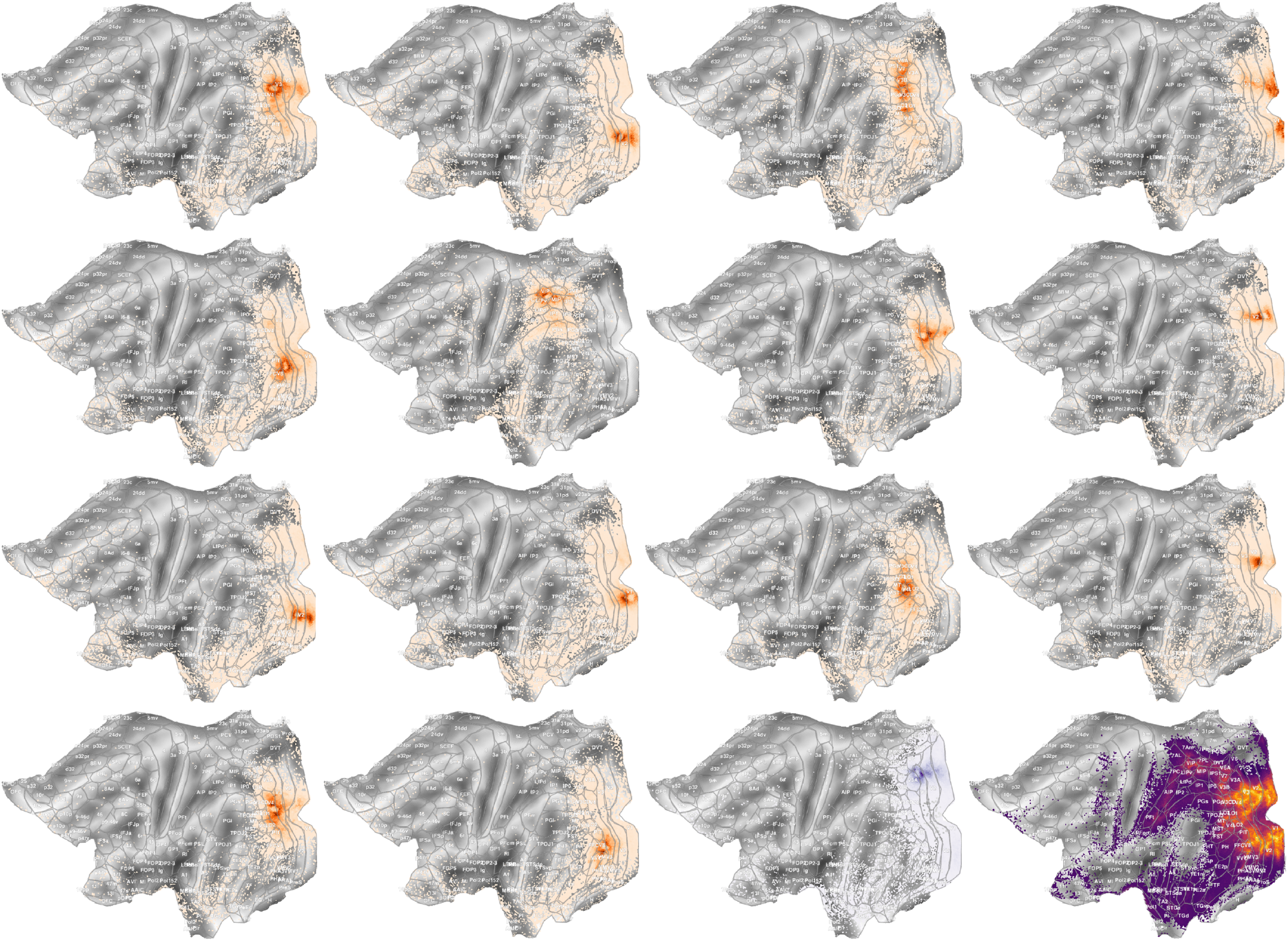
Sparse Dictionary Atoms Derived from Structural Connectivity. Shown are the first 15 atoms from a sparse dictionary decomposition of diffusion MRI-based structural connectivity. Each atom reflects a spatial pattern of influence derived from tractography data. Facilitative atoms (positive weights) are shown in orange, while the suppressive atom (negative weights) is shown in purple. The final panel (bottom right) shows the prediction obtained by linearly combining the selected atoms, representing the model’s reconstruction of retinotopic selectivity. This anatomy-constrained dictionary captures some global features of the selectivity map, but underperforms compared to geometry-derived models such as SCD.

## Data and Code Availability

The datasets used in this study are publicly available at the following sources:

- Retinotopic Selectivity Maps from the Human Connectome Project[16]: https://osf.io/bw9ec/
- Retinotopic Selectivity Maps from the NYU Replication Dataset[22]: https://osf.io/e6vqk/
- Functional Connectivity Gradients [68]: https://www.neuroconnlab.org/data/

The notebooks used to generate all figures in this paper is available at: https://github.com/ulyssek/shaihulud_figures

The implementation of the Spatial Component Decomposition (SCD) algorithm is available at: https://github.com/ulyssek/spatial_component_decomposition

## Acknowledgements

This research was funded by the European Research Council (ERC) under the European Union’s Horizon 2020 research and innovation programme (Grant agreement No. 866533) awarded to D.S.M., and supported by the NIHR Oxford Health Biomedical Research Centre (NIHR203316). The views expressed are those of the author(s) and not necessarily those of the NIHR or the Department of Health and Social Care. The Wellcome Centre for Integrative Neuroimaging was supported by core funding from the Wellcome Trust (203139/Z/16/Z and 203139/A/16/Z).

We thank all members of the CANN lab for their support and feedback, particularly Tsvetoslav G. Ivanov, Eva Sevenster, Aswathi Thrivikraman, Dabal Pedamonti, and Rahul Gupta, for their helpful discussions and insights throughout the project.

The modeling work was carried out using the high-performance computing resources of the Oxford Biomedical Research Computing (BMRC) facility. We are especially grateful to the BMRC support team, and in particular Charles Robert, for his patience, technical expertise, and pedagogical support.

## Author information

Authors contributed equally: S.F.-W., D.S.M.

## Contributions

U.K., P.-L.B., S.F.-W., and D.S.M. conceived the study (Conceptualization) and developed the methodology (Methodology). U.K. implemented the software (Software), performed the analyses (Formal analysis, Investigation), and created the visualizations (Visualization). U.K. also led the initial draft of the manuscript (Writing – Original Draft). All authors contributed to reviewing and editing the manuscript (Writing – Review & Editing). P.-L.B., S.F.-W., and D.S.M. provided supervision and project oversight (Supervision), and D.S.M. acquired funding for the project (Funding acquisition).

## Corresponding author

Correspondence to S.F.-W., D.S.M.

## Competing Interests

P.-L.B. is the owner of Full Brain Picture Analytics. The remaining authors declare no competing interests.

## AI Usage

GitHub Copilot and ChatGPT were used throughout the scientific process. These tools contributed to code review and refactoring, suggested programming architectures and data structures, and assisted in reducing grammatical errors and typographical mistakes, as well as improving the clarity and readability of the manuscript. All AI-generated suggestions were carefully considered and reviewed by the authors.

